# Sexual dimorphism in the dorsal spot number and yellow surface in fire salamanders *Salamandra salamandra ssp terrestris* Linnaeus, 1758 (Caudata: Salamandridae)

**DOI:** 10.1101/2024.03.25.586466

**Authors:** Martin Bozon, Basile Marteau

## Abstract

Sexual selection among amphibians is mainly based on size and colour dimorphism. Those criteria are less studied in salamanders. Only morphometric and yellow surface differences between males and females are known. We studied a *Salamandra salamandra terrestris* population and observed significant differences in spots number and yellow surface area between males and females. Females have on average more spots and are blacker than males on their back.

---

Sexual selection is one of the primary factors that can explain the morphological diversity of living species. Proposed by Darwin in 1871, the study of this evolutionary process continues because it allows us to understand how a species interacts with its environment and can help to explain its development, its evolution or the way the species adapts to its environment (Servedio and Boughman, 2017).

Amphibians exhibit a wide range of sexual selection criteria such as their body size, song, colours and the use of pheromones (Andersson, 2019). In many species of amphibians, colour is a signal that varies temporally, appearing and disappearing during the reproductive period (Bell and Zamundio, 2012). Pigmentation can increase individual reproductive success influenced by courtship, partner recognition or fitness perception (Davis and Grayson, 2008; Andersson, 2019). This phenomenon was demonstrated in the literature for several species of anurans such as the moor frog, *Rana arvalis* (Sztatecsny et al., 2012). However, very little studies have been conducted in urodeles. For example, it is accepted that for some newt species of the genus *Triturus sp.,* sexual selection is based on pheromones, time of the courtship or caudal filament length (Cornuau et al., 2012). The role of the coloration has been suspected but never proven. The most recent study was done by Secondi et al., (2012) on the Smooth newt, *Lissotriton vulgaris* showing that females tend to spend more time near males who reflect more UV light on the ventral side.

In some amphibian species such as the *Dendrobatidae* family, it has been shown that the colour display by individuals is an aposematic signal (see review Santos et al., 2016). Similarly, the yellow colour of the fire salamander has always been considered as an aposematic pattern (Parker and Bellairs, 1971; Seidel and Gerhardt, 2016). In fact, the venom secreted by the fire salamanders is extremely toxic and deadly to its predators and to the salamanders themselves (Rollard et al., 2015). Except for some rare cases where salamanders have been predated by rats (Velo-Antón et al., 2011), fire salamanders have no predators in their adult stage. However, in their review, Lüddecke et al., (2018) question this uniquely aposematic function.

The fire salamander, *Salamandra salamandra*, is a common urodele distributed throughout the European continent (Thornn and Raffaelli, 2001). Due to this wide distribution, many morphological and behavioural variations exist. In France, its observation remains occasional due to its nocturnal and underground habits (Speybroeck et al., 2018). The adult fire salamander measures between 14 and 20 cm and displays yellow dorsal spots patterns. These colour patterns are specific and unique to each individual, varying in terms of number and shape of the yellow spots (Drechsler et al., 2014). Sexual dimorphism is not marked in this species, and sex identification is possible based on the size of the cloaca during the reproductive period (Thornn and Raffaelli 2001).

The mating and reproduction processes of the fire salamander are widely studied. Many criteria such as size (Labus et al., 2013), olfaction and pheromone emission (Caspers and Steinfartz, 2011) have been been identified as sexual selection criteria. Recent studies showed that fire salamanders’ spots are not just an aposematic sign, but is also sexually dimorphic (Balogová et al., 2015; Preißler et al., 2019) which is manifested not by a difference in the number of spots, but by the total area of the yellow colour on the salamander’s body.

The aim of this study is to investigate the difference in dorsal spot number and yellow surface between males and females of the fire salamander, *Salamandra salamandra terrestris.* We expected to find similar results as Balogová et al. (2015) and Preißler et al. (2019). The goal of those two papers were different, Balogová et al. (2015) focused on the sexual dimorphism of spot number and yellow ratio on the back and on the front legs of *Salamandra salamandra* ssp. *salamandra* while Preißler et al. (2019) focused on the relationship between yellow ratio and toxicity and noticed a link between yellow ratio and sex amongst *S. salamandra* ssp. *terrestris*. Both papers found yellower males compared with females but none of them focused or found any significant difference of spot number between sexes. We expected to observe more yellow spots within the female group. If the females are blacker than males, then the number of intersections between spots is higher and so the black surface.

## Material and methods

### Study Area and Protocol

We conducted the study in the Parc de Saint-Nicolas, in Angers, France. All the pictures were taken during a Capture-Mark-Recapture protocol composed of 6 transects (Fig. 1) surveyed for 3 nights (10/10/2021, 02/11/2021 and 21/11/2021). Salamanders’ dorsal patterns being unique (Kiss et al., 2021), the protocol is based on individual picture recognition and didn’t necessitate any manipulations. The picture data bank is composed of almost 1000 individuals.

**Figure 1:**
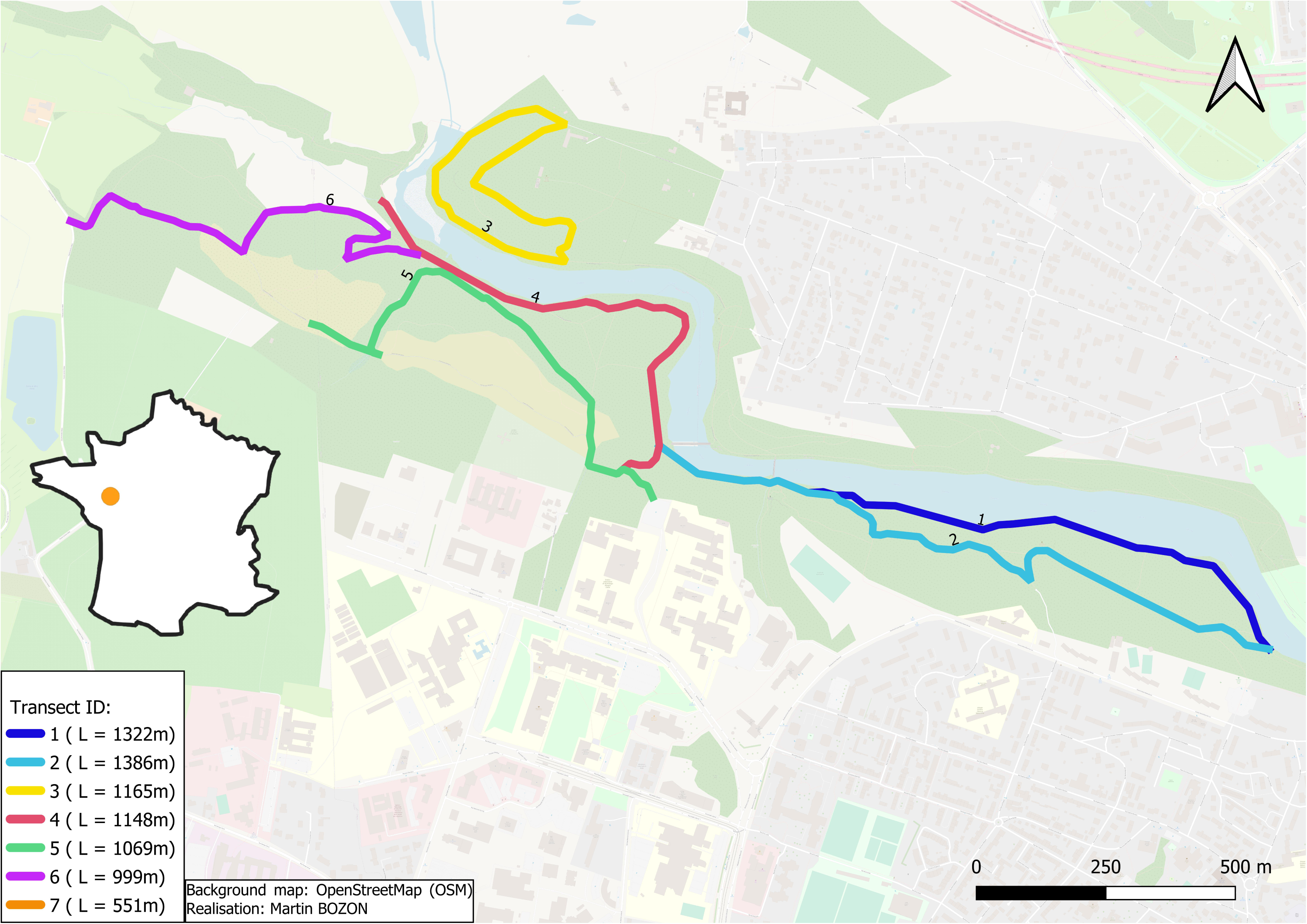
Prospected transects during CMR sessions, L= length in meters.

Pictures were taken with different cameras and smartphones from several brands.

### Sex Determination

Sex determination was made in the field and also later with pictures taken by PEGAZH non-profit organisation members.

During the breeding season, males’ cloaca is swollen and stand out well from the tail (Fig. 2 A; Thornn and Raffaelli, 2001) whereas females’ cloaca stays flat. Pregnant females are easily recognisable due to their bloated belly (Fig. 2 B). Only individuals that could be sexed were taken into account in the analyses.

**Figure 2:**
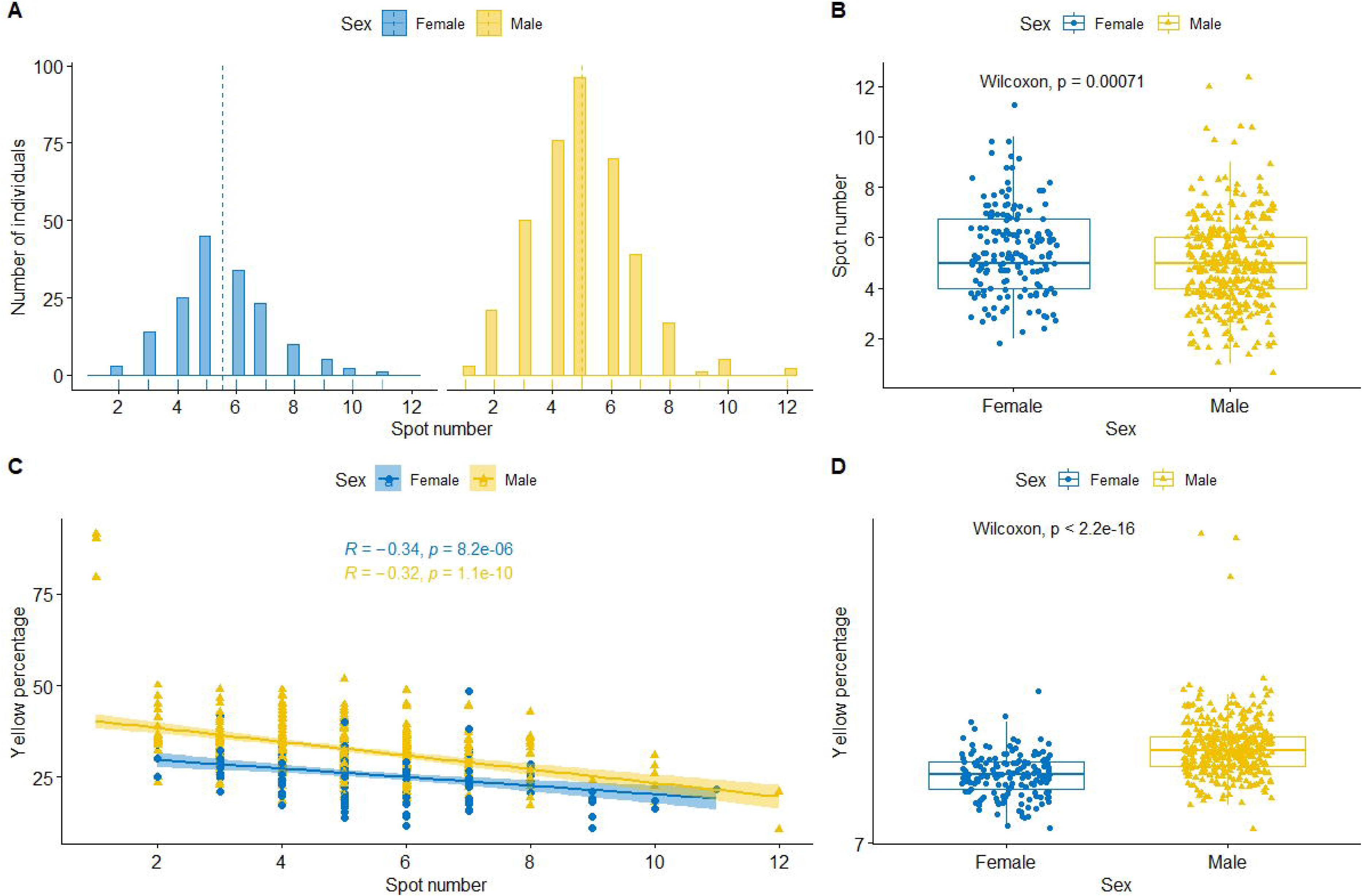
A) Male with swollen cloaca; B) Pregnant female.

### Spot Number and area Determination

Spot number corresponds to the number of spots included along the two dorsal longitudinal lines (specific to the *terrestris* subspecies) inside the area located between the base of the neck and behind the hind legs (Fig. 3 A). Those dorsal spots are easily identifiable and very little counting mistakes can be done.

**Figure 3:**
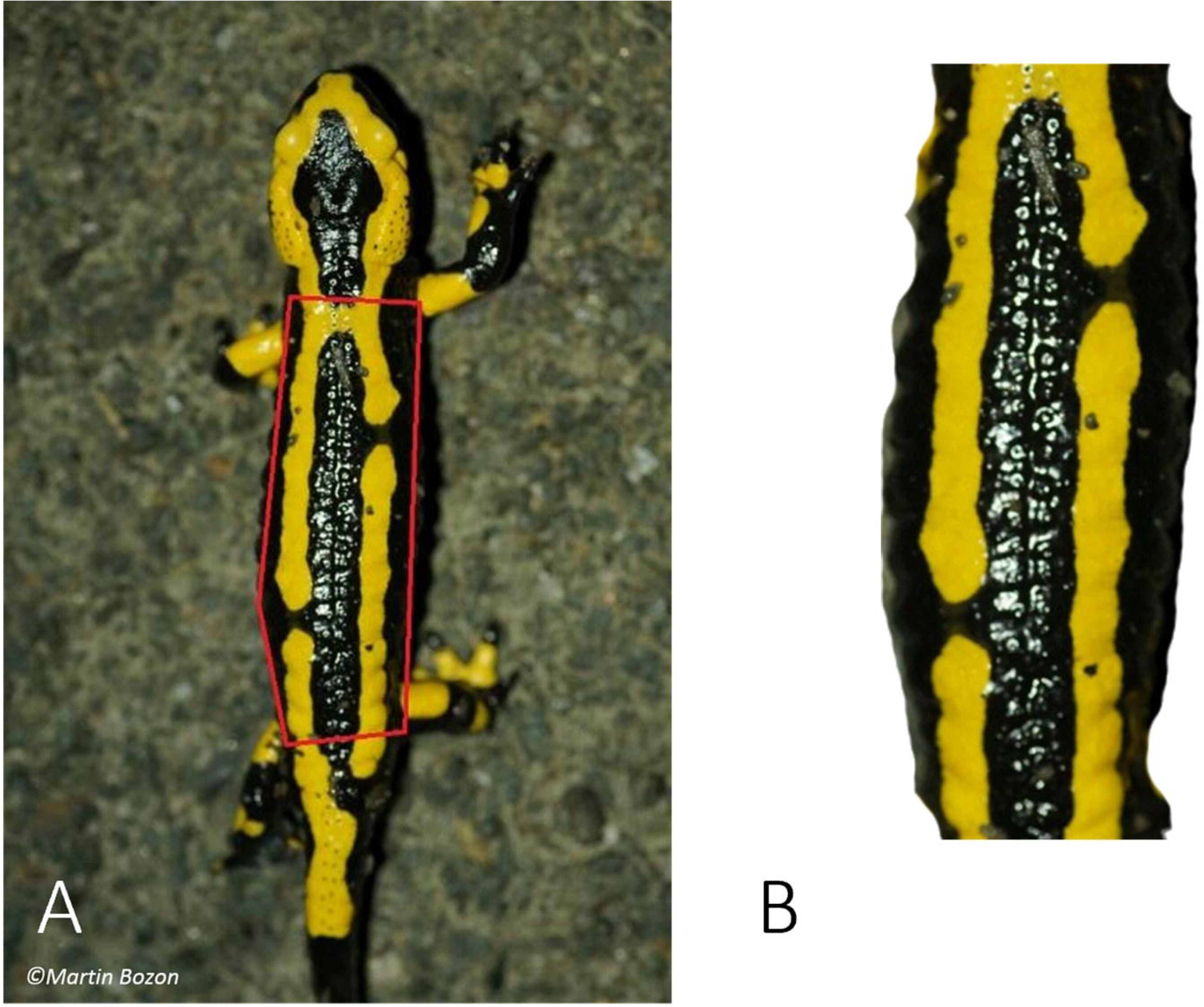
A) Spot counting area; B) area for yellow surface calculation.

The surface area of yellow pigmented skin was also estimated. using the Python algorithm developed by Sanchez et al., (2018) that calculates the percentage of yellow pixels in the pictures. Because the pictures were not standardised, cropped the pictures manually (Fig. 3 A).

### Statistical Analysis

We performed all statistical analysis under R using the Rstudio environment (Rstudio version 1.4.1717).

The spot numbers and the yellow percentage were not normally distributed (Shapiro test; *P*-value < 0.01). Consequently, The R package “MASS” (Venables and Ripley, 2002) was used to run generalised linear models (GLM). We applied a backward selection process using the “drop1” function with a Chi test. Normality of residuals and homoscedasticity of the models were analysed with the “DHARMA” package (Hartig, 2016).

The sex effect on yellow percentage and the number of dorsal spots were assessed with a GLM. Finally, the relationship between the number of spots and the yellow percentage was investigated with a Pearson correlation. In the two models, GLM had a gaussian family distribution and a logit link function. The yellow percentage was log transformed to meet the criteria of residual normality.

## Results

In total, 553 salamanders were sexed, 389 males and 164 females (Table I). During the fieldwork complete yellow males were observed. So, the dorsal spot number for males went from 1 to 12 and for females from 2 to 11. This yellow morph is relatively common for *ssp terrestris* (Rimpp, 1984; Thornn and Raffaelli, 2001). No females of this specific pattern were found during the survey. The yellow percentage extends to a range from 91.35 (yellow male individual) to 11.22 (a female).

**Table I:**
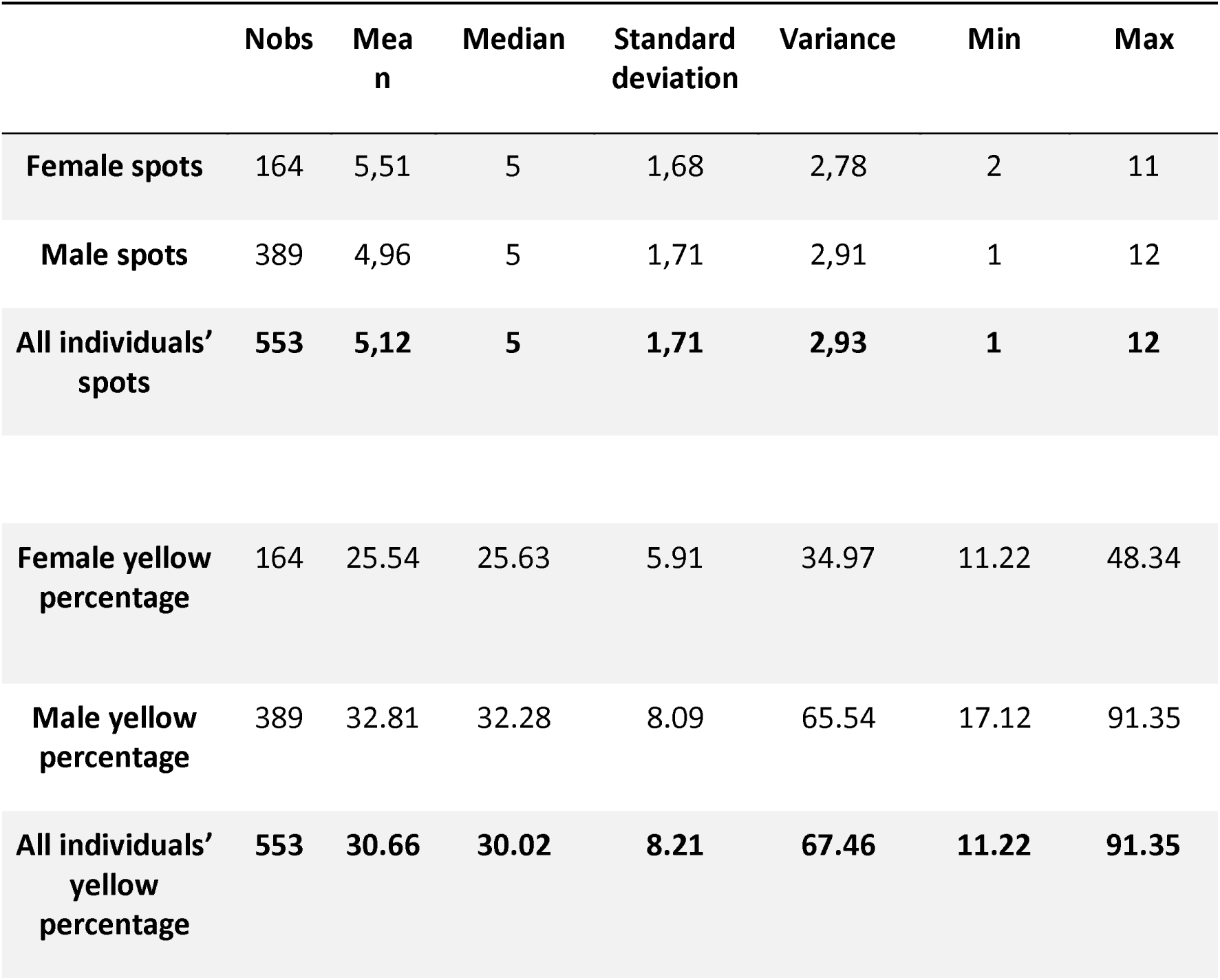
Summary of the results on the number of spots between the sexes and the yellow percentage difference between sexes for the Salamander population present in the study area.

|  | **Nobs** | **Mean** | **Median** | **Standard deviation** | **Variance** | **Min** | **Max** |
| --- | --- | --- | --- | --- | --- | --- | --- |
| **Female spots** | 164 | 5,51 | 5 | 1,68 | 2,78 | 2 | 11 |
| **Male spots** | 389 | 4,96 | 5 | 1,71 | 2,91 | 1 | 12 |
| **All individuals' spots** | **553** | **5,12** | **5** | **1,71** | **2,93** | **1** | **12** |
| **Female yellow percentage** | 164 | 25.54 | 25.63 | 5.91 | 34.97 | 11.22 | 48.34 |
| **Male yellow percentage** | 389 | 32.81 | 32.28 | 8.09 | 65.54 | 17.12 | 91.35 |
| **All individuals' yellow percentage** | **553** | **30.66** | **30.02** | **8.21** | **67.46** | **11.22** | **91.35** |

**Table II:** Results of the GLM performed between the sex and the spot number, between the sex and the log of the yellow percentage.

| Model type | Response variable | Explanatory variable | Model results |  |  |  |
| --- | --- | --- | --- | --- | --- | --- |
| GLM<br>(Gaussian family) | Spot number | Sex | Fixed effects | Estimate | Standard deviation | *P-value* |
|  |  |  | Intercept | 5,51 | 0,13 | < 0,001 |
|  |  |  | Male | -0,55 | 0,16 | < 0,001 |
| GLM<br>(Gaussian family) | Log(Yellow percentage ) | Sex | Fixed effects | Estimate | Standard deviation | *P-value* |
|  |  |  | Intercept | 3,21 | 0,02 | < 0,001 |
|  |  |  | Male | 0,25 | 0,02 | < 0,001 |

The model selection using a Chi test revealed a significant effect of sex on the spot number (*P*-value < 0.001) and on the yellow percentage (*P*-value <0.001; table I). The GLM (Gaussian law, link function logit) shows males having significantly fewer spots than females (intercept= 5.51, males= –0.55; Figure 4 A and 5 B). Models also revealed a significant sex effect on the yellow percentage; males are significantly yellower than females (intercept= 3.21, males= 0.25, Figure 4 D). We also demonstrate that the number of spots is correlated with the yellow percentage. The fewer spots there are, the yellower the salamanders are (males, r= –0.32, *P*-value < 0.001; females, r=-0.34, *P*-value < 0.001, Figure 4 C).

**Figure 4:**
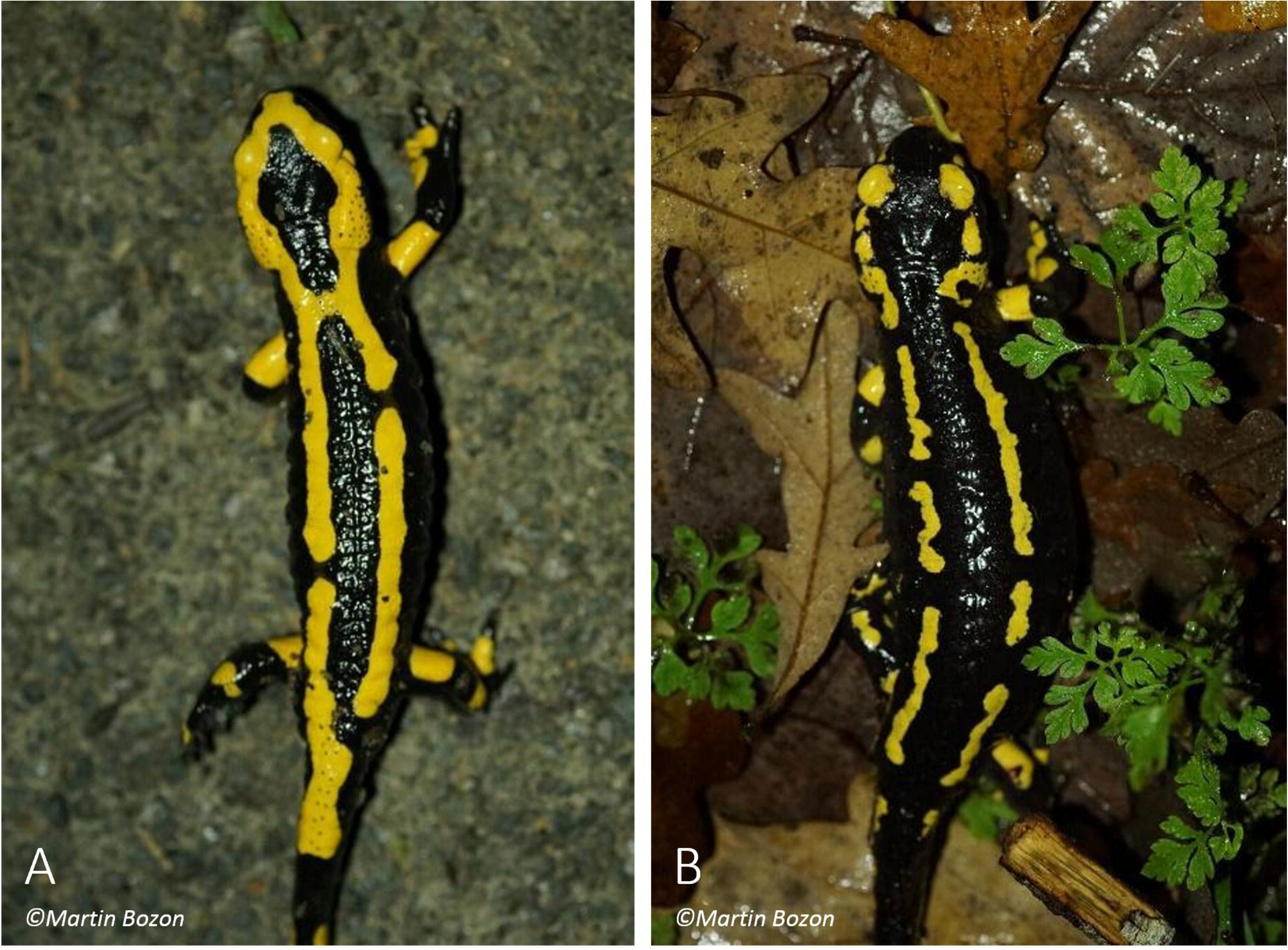
A) Spot number distribution for males and females, dotted lines show the mean number of spots for each sex; B) Boxplot of the difference of spot number between males and females (Wilcoxon test, P-value < 0.001); C) Correlation (Pearson) between the spot number and the yellow percentage (males: r= –0.32, P-value < 0.001; females: r= –0.34, P-value <0.001); D) Boxplot of the difference of yellow percentage between males and females (Wilcoxon test, P-value < 0.001.

## Discussion

This study revealed a significant difference in the number of dorsal spots, females have on average more spots than males in this population of fire salamanders. Dorsal yellow percentage differences were also detected between the sexes showing males being yellower than females. Our test also revealed a significant link between the spot number and the yellow percentage. The less there are dorsal yellow spots, the more yellow the salamander is.

Our study’s results agree at least partly with those of Balogová et al., (2015) where they demonstrated that the males, compared to females, were more yellow on the back and the tail. However, they did not find any dorsal spot difference between the sexes. We believe the absence of spot number differences in Balogová et al., (2015) may be explained in two ways. Firstly, the difference in the sub-species between the studies may explain the difference. The yellow spot patterns differ greatly between the two subspecies. *Salamandra s. salamandra* tends to have an irregular pattern of spots that are sometimes arranged in longitudinal bands while *Salamandra s. terrestris* spots are arranged in two yellow stripes along the back (Rimpp, 1984; Thornn and Raffaelli, 2001; Sparreboom, 2014).

Secondly, the small number of individuals (n=68) in Balogová et al., (2015) from different populations may not be enough to detect a significant difference in the number of spots between males and females.

The yellow pattern function of the fire salamander remains unclear. Preißler et al., (2019) demonstrated that the yellow surface area is not representative of the toxicity level of the individual. The main use of this yellow dorsal pattern was only described as an aposematic trait (Parker and Bellairs, 1971; Seidel and Gerhardt, 2016). However, many authors doubt this function. For example, Preißler et al., (2019) and Rivera et al., (2014) hypothesise that yellow colour could be mainly used as camouflage on the forest substrate during fall and is also a sexually dimorphic trait showing the individual quality, and so be an honest signal (Zahavi, 1975). In France, salamanders’ peak activity is in autumn and winter (Thornn and Raffaelli, 2001). Then, the yellow colour could help to hide between the leaves and lower the risk of predation.

The differences in dorsal yellow percentage per sex coupled with the differences of dorsal spots numbers could open the possibility that the yellow colour is used in courtship display by the salamanders (Balogová et al., 2015; Preißler et al., 2019). After the male’s courtship, where it goes underneath the female to mate, she could decide to accept the spermatophore or not depending on the male’s back coloration. Recent studies show that the yellow percentage in fire salamanders is directly linked to the food availability during larval stage (Barzaghi et al., 2022). Larva that has grown in a more nutrient-rich environment will show a higher yellow percentage after the metamorphosis. The yellow pigmentation is caused by a high concentration of carotenoid in the salamander’s skin (Lucchini and Pizzigalli, 2020). This family of pigment is commonly used in the immune response (Chew, 1993; Chew and Park, 2004). Exhibiting flashy patterns mainly composed by carotenoids is an honest signal for potential mates to attest to the potential of the individual health quality (Zahavi, 1975). Lots of animal species like birds or mammals use such signals during sexual courtship to seduce females (Pike et al., 2007). Like other amphibians (Rudh and Qvarnström, 2013), the male’s coloration could be indicative of an individual’s quality, helping females to select a mate.

The similar results of the yellow surface difference between males and females found by Balogová et al., (2015) and Preißler et al., (2019) suggests that sexual dimorphism in pigmentation may also exists in other subspecies of fire salamanders, and maybe other salamanders species. The strong and significant correlation between the spot number and the yellow surface prone that the yellow coloration, and so spot number, could be a sexually dimorphic trait in the fire salamander.

This question deserves further investigation. If similar results came to be observed, a laboratory experiment based on mate selection by females could be led to determine if the difference in yellow surface, and so the number of spots comes from female selection or if other factors are at play such as the testosterone level. We believe that such studies will deepen the knowledge on this common yet little known and threatened urodele.

## Acknowledgments

We would like to thank all the PEGAZH non-profit organisation members who participated in the survey, all of this wouldn’t have been possible without them. We also want tomakea special thank to Leo SEPTIER for the time he spends helping to find money. And finally, we want to thank Corentin PAPE and John LOEHR for proofreading and their advice.

